# Land use change drives major loss of Southeast Asian biodiversity

**DOI:** 10.1101/2023.08.08.552370

**Authors:** Thomas Botterill-James, Luke A. Yates, Jessie C. Buettel, Zach Aandahl, Barry W. Brook

## Abstract

Southeast Asia is highly biodiverse and currently experiences among the highest rates of tropical deforestation globally, but impacts on biodiversity are not well synthesized. We use Bayesian multi-level modeling to meta-analyse 831 pairwise comparisons of biodiversity in sites subject to human land use change and anthropogenic forest disturbance (for example in plantations or logged forest) versus undisturbed sites. After controlling for hierarchical dependences, we show that biodiversity is a fifth lower in sites with these land-use changes (95% credible interval= 16-28%, mean = 22%). This reduction was greater when land use change/anthropogenic forest disturbances were high-intensity (34% reduction in biodiversity) compared to low-intensity (18% reduction), and effects were consistent across biogeographic regions and taxa. Oil-palm plantations lead to the greatest reduction in biodiversity (39%, CI 27-48%), and agroforests the least (24%, CI 10-37%). We also find that biodiversity is reduced in young secondary forest by 26% (CI 4-42%) compared to undisturbed forest, but there is no reduction in biodiversity for intermediate or mature-aged secondary forest (although species composition is potentially altered). Overall, our study provides the clearest evidence yet of the substantial detrimental impact of land-use change and anthropogenic forest disturbance on the biodiversity of Southeast Asia.

## Introduction

A substantial fraction of future extinctions caused by anthropogenic global change are expected to occur in tropical forests because the Earth’s terrestrial biodiversity is concentrated within these ecosystems^1-3^. Despite covering less than 10% of the Earth’s surface^4^, tropical forests are estimated to support at least two-thirds of the world’s terrestrial biodiversity^5,6^. This biome is threatened by many chronic human-mediated pressures, including hunting^7-10^, the wildlife trade^11-13^, climate change^4,5^ and habitat degradation and fragmentation (e.g., from intrusion of logging roads^14,15^). However, the greatest threat to biodiversity in these ecosystems comes from large-scale land-use changes and anthropogenic forest disturbances (interchangeably used here to describe any major human use of forested areas that results in deforestation of previously undisturbed or minimally disturbed forest)^16-19^. Over half of the original extent of tropical forests globally have been lost in modern times, likely causing or committing some 15% of tropical forest species to become extinct due to direct or time-lagged effects^20^. Tropical biodiversity is expected to come under even greater pressure into the future, as increasing global human population and associated demand for agriculture and other commodities, including from post-forest transition areas (e.g., China, India, G7 nations) drives expanding deforestation^21-28^. Moreover, pressure on biodiversity in tropical areas will be amplified compared to non-tropical regions because: 1) much of the forecast growth in global population will occur in the tropics, 2) these regions are experiencing marked increases in living standards and per-capita natural resource consumption, and 3) deforestation for agricultural and bioenergy production is likely to be higher in the tropics, with cheap land and tropical climates facilitating a year-round growing season^26^.

Nearly 15% of the area of the world’s tropical forests are in Southeast Asia^29^ and the region is recognized for its disproportionately high endemism at the species and higher taxonomic levels^30,31^. Unfortunately, Southeast Asia is also a deforestation hotspot, accounting for a large amount of the ongoing deforestation occurring in tropical humid and lowland forests^30,32-36^. This translates to an elevated threat relative to other regions: compared with Meso-America, South America and sub-Saharan Africa, Southeast Asia has the highest regional proportion of vascular plant, reptile, bird, and mammal species classified as globally threatened on the IUCN’s Red List^30^. Further, it is forecast that up to half of Southeast Asia’s terrestrial species could be regionally extinct by 2100 if currently modelled rates of habitat loss continue^35,37^. As such, understanding land-use-change effects on biodiversity in this region is a crucial and high-priority component of conserving global biodiversity.

There have been several important recent efforts to synthesise land-use-change effects on biodiversity from a planetary perspective^38-43^. Yet biodiversity responses to land-use change can vary greatly across regions and ecosystems^44,45^. Global approaches, in their overarching scope, are likely to miss important geographical variation in the sensitivity of biota to land-use change^46^ or potentially even mischaracterise responses in some regions. For example, agricultural fields decrease biodiversity in tropical areas^46-49^ but increase biodiversity in desert regions^50,51^. Clearly, more geographically focused approaches are warranted for an improved understanding of land-use change impacts on biodiversity at the regional scale. This will better guide the prioritisation and implementation of conservation decisions on the scale at which action often occurs^44^.

Southeast Asia is a dynamic tropical region undergoing rapid land-use changes^34,52^, but it has been over ten years since responses of biodiversity to human impacts have been synthesised and investigated^30,48^. Further, past syntheses relied on quantitative methods that are now outdated (for example, unweighted meta-analysis). Here, for the first time, we comprehensively quantify and explore land-use-change impacts on biodiversity for this region, using location-scale meta-analytic methods with inverse-variance weighting^53-56^. We undertake a comprehensive and systematic review focused on Southeast Asia for the decade spanning 2010–2019, collecting 831 pairwise comparisons (from 101 studies) of biodiversity values in primary or mature secondary forest (hereafter undisturbed) to areas affected by land-use change and anthropogenic forest disturbance (Fig. 1). We use Bayesian multi-level models to meta-analyse the standardised mean difference between undisturbed and human-affected areas, quantifying the overall change in biodiversity between them whilst controlling appropriately for hierarchical relationships (in particular, by accounting for non-independent observations caused by structured inter-dependencies). We then use a suitable form of cross-validation to compare a set of candidate models varying in their combination of predictors, to understand which components are most important for moderating the effects of land use change on biodiversity. We focus on three principal predictors: intensity of land use change/disturbance intensity (high or low, classification detailed in material and methods), biodiversity metric (species richness, abundance, or forest structure), and taxonomic group (plant, invertebrate, or vertebrate). We also calculate log-response ratios from both intercept-only (null) and best-supported models, so that biodiversity change is expressed throughout as an easily interpretable percentage change (with 95% credible intervals to show uncertainty around estimates).

**Figure 1.**
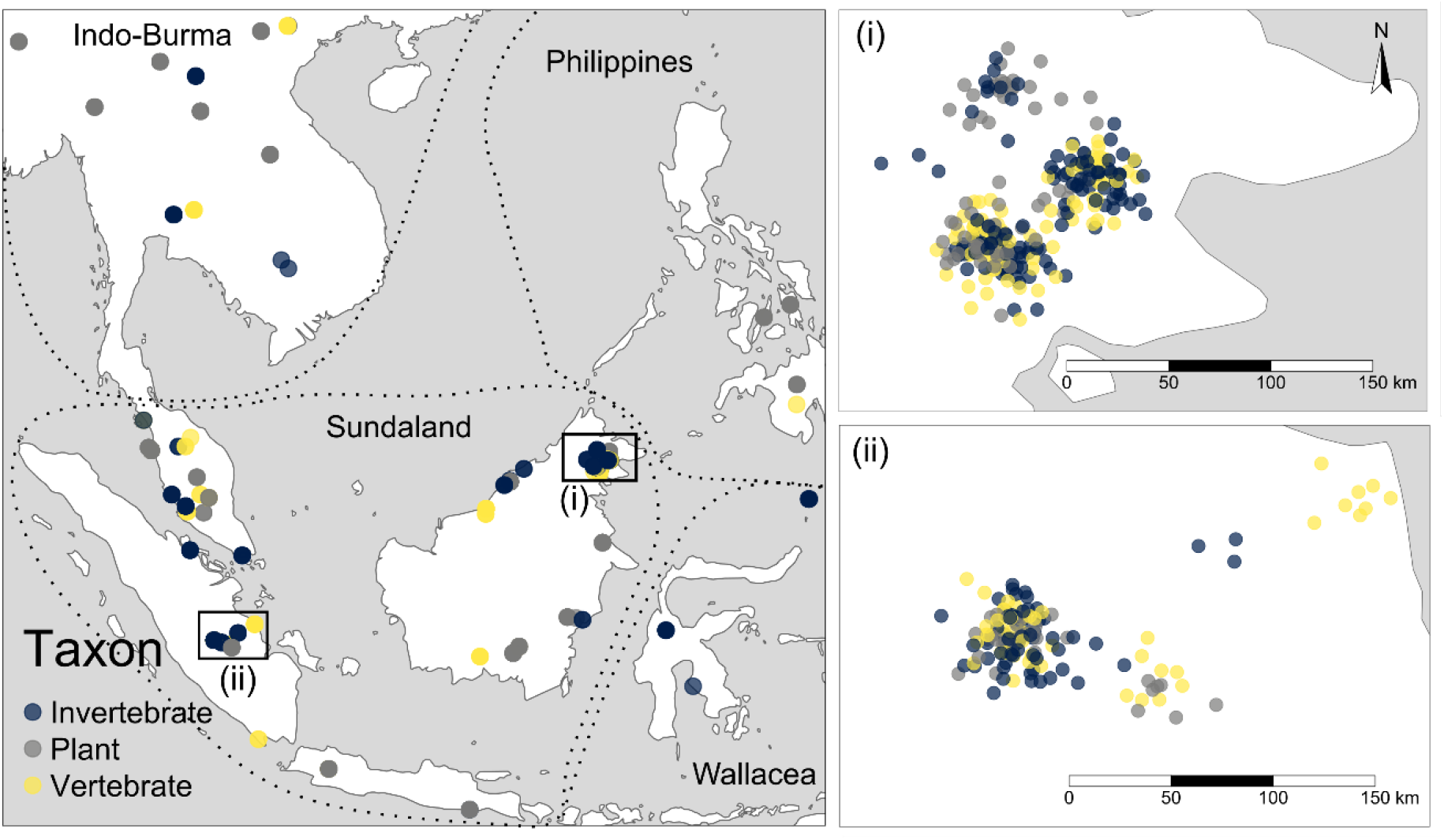
Map of studies (n = 101 studies, and 831 effect sizes) included in the meta-analysis, with dashed lines demarcating major biogeographic regions. Studies are grouped by taxon category. *Zoomed insets (i) and (ii)* show details for areas with the highest clustering of samples, being: (i) the Indonesian province of Jambi, and (ii) the Malaysian province of Sabah. Points are randomly jittered by ∼5km to reduce overlap, and so represent approximate locations.

## Results and Discussion

The intercept-only model (comparing only control versus impact sites, irrespective of intensity, metric or taxon) showed that there was a marked decrease (22%, CI = 16-28%) in biodiversity in human-affected areas. This effect is stronger than that found in a recent global synthesis quantifying land-use effects on vertebrate biodiversity (11.1%)^41^, but comparable to a previous analysis focused on biodiversity decline in Southeast Asia between 1975 and 2007 (where biodiversity was 22.2% lower in disturbed compared to pristine sites)^48^.

Although our intercept-only model revealed an overall decrease of biodiversity, there was substantial unmodelled variation in effect size between studies, as measured by the *I*^2^ statistic. *I*^2^_total_ – the total proportion of effect-size variance, ignoring the total known sampling variance^57^ – was 37% (CI 27-48%), which is lower than typically high heterogeneity often found in ecological datasets^58^ due to our use of location-scale models which predict both the mean and variance of the response variable^54^.

We then explored the potential causes of this heterogeneity, performing model selection on nine candidate models representing varying combinations of the predictors (disturbance intensity, biodiversity metric, and taxon) and their interactions, with the candidate model set including a null, intercept-only model. Based on 20-fold leave-cluster-out cross validation (see material and methods for model selection details), the model that most parsimoniously fit the data included both disturbance intensity and biodiversity metric as predictors but left out the taxonomic breakdown (Fig. 2).

**Figure 2.**
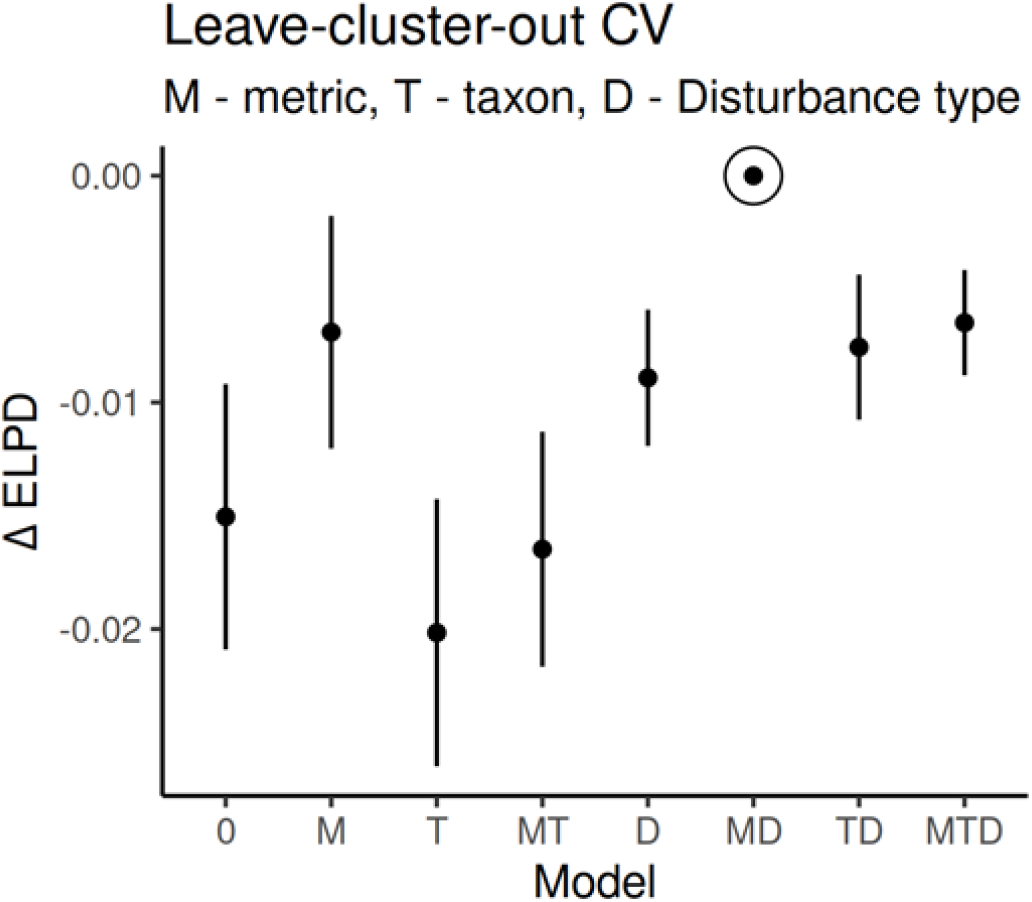
Relative performance of models of biodiversity metric, taxon and disturbance intensity effects and their interactions on biodiversity loss in Southeast Asia. Out-of-sample model performance was estimated using 20-fold leave-one-group(cluster)-out cross validation. Models are grouped on the x-axis according to the number of fixed effects, increasing in complexity from left to right. Model performance was quantified using expected log-pointwise predictive density (ELPD), expressed here as the relative estimates (Δ ELPD) with respect to the best-performing model. The error bars depict one standard error of the estimated differences, with the performance of an alternative model deemed comparable to the best model if the mean estimate (filled circle) lies within approximately one standard error of Δ ELPD = 0. The best out-of-sample performing model included disturbance intensity and metric type as fixed effects (circled on figure). We therefore use this model for inference throughout the paper, except where we present analyses of disaggregated land-use/disturbance categories on subsets of data.

Increased intensity of land use and anthropogenic forest disturbance has been found to negatively influence biodiversity across a wide range of habitats and biomes^45,59,60^, but previous studies that focused on tropical forests in Southeast Asia failed to detect any moderating effect of intensity on biodiversity^48^. We were able to provide clear evidence that the loss of biodiversity in higher intensity disturbed areas was much greater relative to less-intensely disturbed areas, demonstrating that intensification of land use change and forest disturbance constitutes a strong driver of biodiversity loss in Southeast Asia. Biodiversity was reduced by 35% (CI 27-42%) in high intensity disturbed areas compared to undisturbed controls, and by 18% (CI 11-24%) in low intensity disturbed areas compared to undisturbed controls. These effects were consistent across taxa and are very similar in magnitude to those previously reported in a study of pan-tropical bird biodiversity^61^. Our results also show that while there was a large amount of variation in effect sizes overall, and some individual instances where biodiversity was higher in disturbed compared to undisturbed sites, the overwhelming pattern was for biodiversity reductions when disturbance intensity was high (Fig. 3).

**Figure 3.**
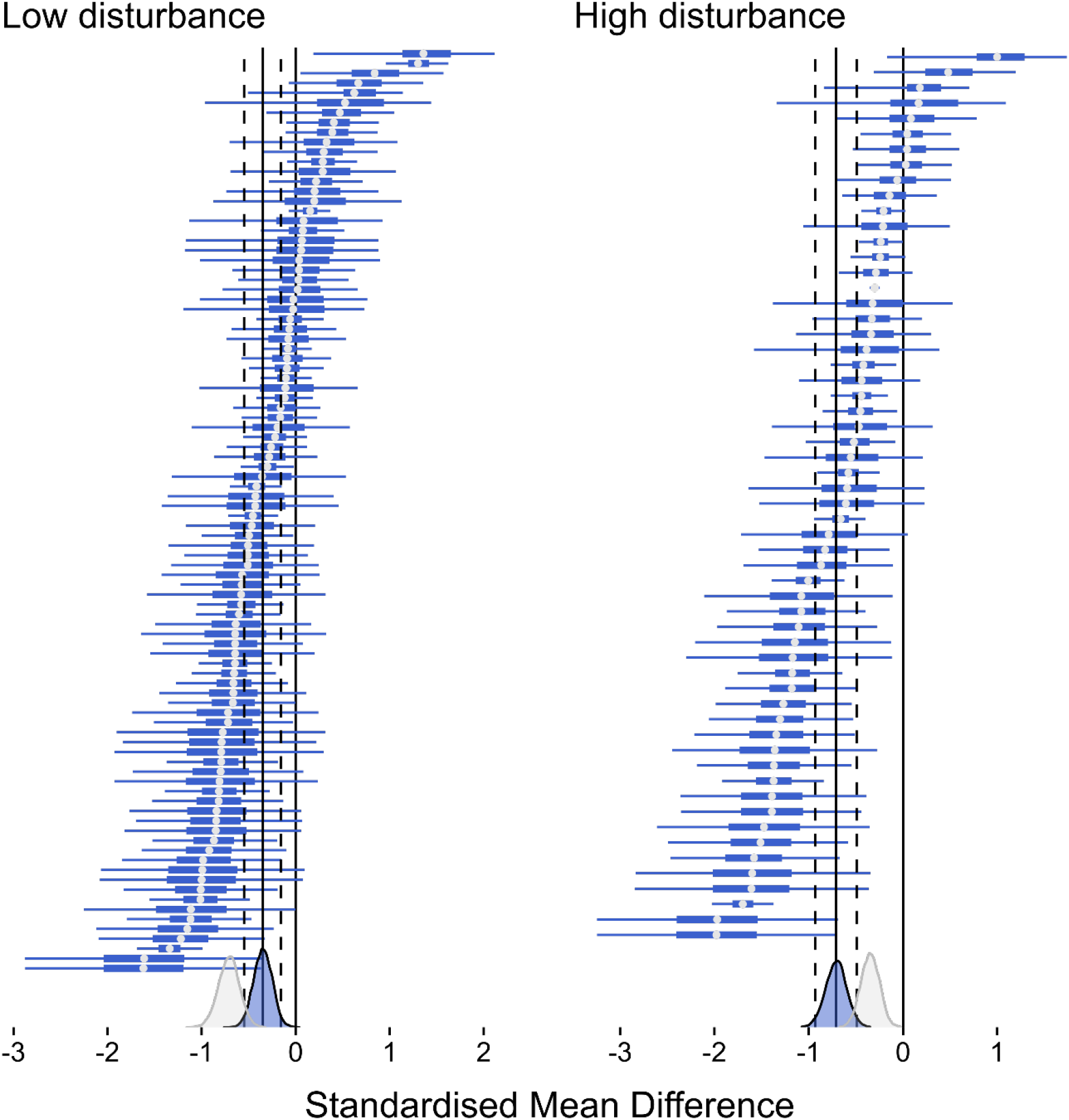
Distribution of effect sizes based on pairwise comparisons of disturbed vs. undisturbed reference sites (represented by the vertical line at 0), under low- and high-intensity disturbances. Points represent the model-predicted mean for each study, with thick and thin bars representing 50 and 95 % credible intervals, respectively. Negative values represent reduced biodiversity in disturbed sites compared to undisturbed sites (the expected direction of impact). At the bottom of each panel, the posterior distribution is shown in blue fill, with the solid line representing the posterior mean and dashed lines representing 95 % credible intervals. The posterior distribution of the other disturbance category is shown in clear fill to facilitate comparison of effect between disturbance intensities.

We also disaggregated our ‘high’ and ‘low’ intensity classifications into finer categories of land-use change/disturbance and fitted the model to three subsets of the data with the log-response ratio as the effect size. We first examined differences between types of plantations, from a subset of 59 studies and 346 effect sizes. This revealed that oil-palm plantations led to the greatest reduction in biodiversity (39%, CI 27-48%), and agroforests the least (24%, CI 10-37%) (Fig. 4A). This finding corroborates results from previous studies investigating the impacts of different plantations on biodiversity^62-64^. The reduced biodiversity impacts of agroforestry compared to monoculture plantations is encouraging, particularly in the context of certain systems such as rubber and cacao where it has been shown that agroforest yields can match those of monocultures^65,66^. However, our results show that biodiversity is still substantially reduced in agroforests compared to undisturbed forests, suggesting that maintaining biodiversity will require ongoing protection of undisturbed forest on a broad landscape scale^48, 67^.

**Figure 4.**
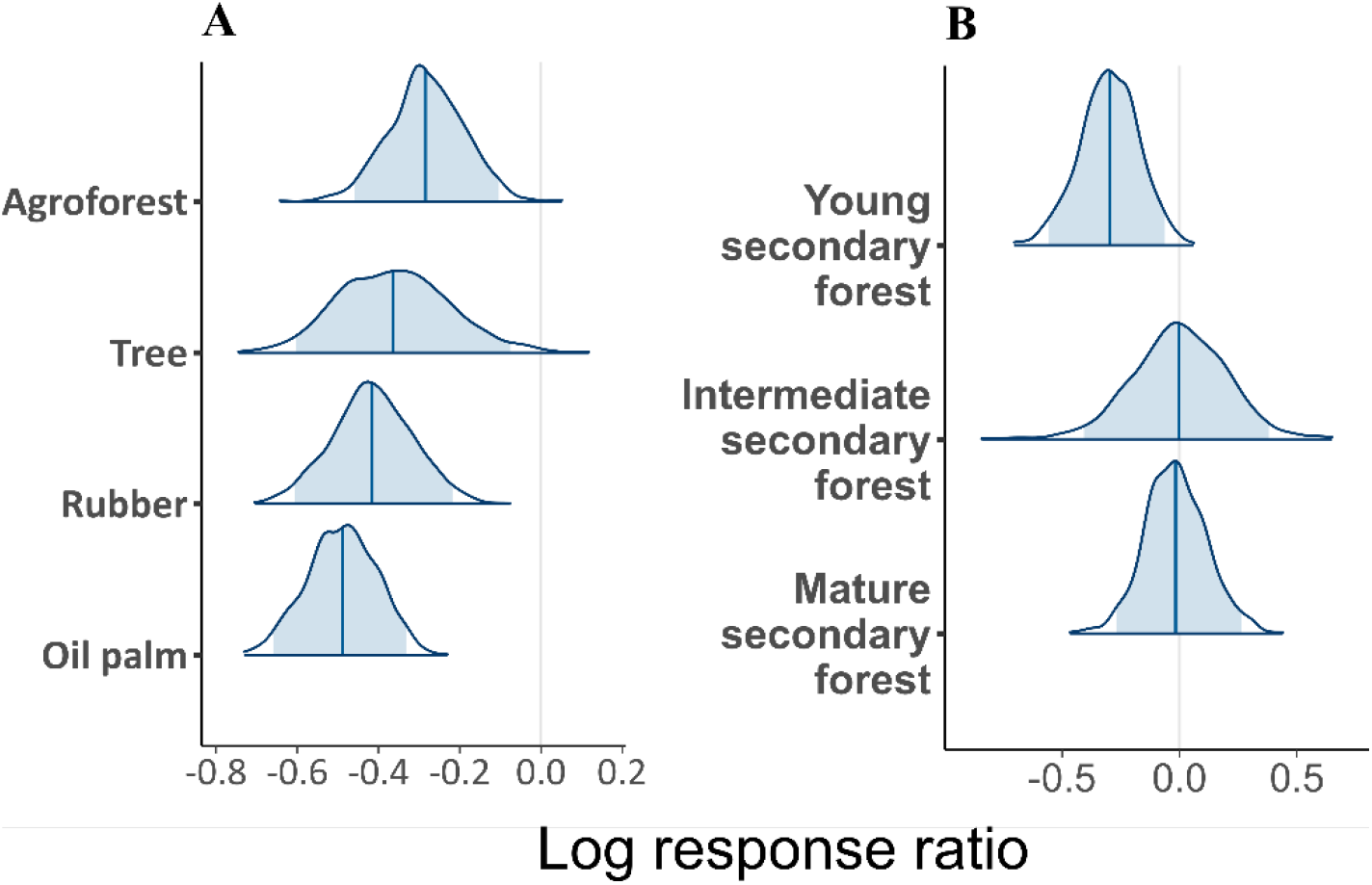
Posterior distributions from **(A)** A model comparing biodiversity in four types of plantations (agroforest, trees – for example, eucalyptus, pine, teak, mahogany, acacia –, rubber, and oil palm) to undisturbed reference sites (represented by the vertical line at 0), and **(B)** A model comparing biodiversity in three ages of secondary forest (young, intermediate, and mature) to undisturbed reference sites. Vertical lines within the distributions show the mean from the model fit for each parameter, with shaded areas representing 95% credible intervals. Negative values represent greater biodiversity reductions in the disturbed sites compared to undisturbed sites (the expected direction of impact).

We then sought to compare biodiversity in mature plantations (both rubber and oil-palm) with undisturbed controls using a different subset of data and models, as it has been suggested that the development of a stable microclimate and complex understorey in these plantations can support diverse communities spanning a range of taxa^68,69^. From our subset of 13 studies and 30 effect sizes, we found that biodiversity in mature plantations was only reduced by 12% (CI: 45% decrease –45% increase, i.e., credible intervals overlapping with no effect). It should be noted that there is a large amount of uncertainty in this estimate (driven by heterogenous effect sizes and our small data subset), and that while our result indicates richness and abundance can recover in these mature plantations, other studies have shown the composition of species assemblages can be substantially altered^70^. Further, any biodiversity recovery within mature rubber and oil-palm plantations is likely to lost upon the destructive replanting process^71-74^. Across Southeast Asia, large swathes of early-generation mature oil palms (i.e., oil-palm plantations that replaced forest during the large-scale oil-palm expansion in the 1990s) are now being replanted or are likely to be replanted in the near future^73,74^, presenting a recurrent threat to biodiversity in the region. There is a current gap in understanding of the responses of biodiversity to oil palm replanting (but see^71-75^), and more research is needed to understand the potential consequences of the next cycle of replanting.

Finally, we examined differences between secondary forests at different stages of recovery from disturbance (young, intermediate, or mature, classified in line with the original study classification where provided, or otherwise classified as young: <10 years, intermediate: 10-20 years, and mature: >20 years). From a subset of 20 studies and 108 effect sizes, we found that biodiversity was reduced in young secondary forest by 26% (CI 4-42%) compared to undisturbed forest (Fig. 4B). In contrast, there was no reduction in biodiversity for intermediate or mature-aged secondary forest (Fig. 4B).

Mature secondary forests have been increasingly considered as viable options for biodiversity conservation because of their ability to host old growth species^76,77^, a supposition that is somewhat supported by these results showing that overall number of species is similar in mature secondary forests compared to primary forests. However, retention of remaining primary forests will remain critical for biodiversity conservation^49^, because after initial conversion, secondary tropical forests take decades to recover the same number of species of old-growth forests and centuries to recover species composition, and forest structure that supports forest adapted species (e.g., tree hollows and fallen wood)^43,78^.

Richness was the most reported biodiversity metric (constituting over half of all pairwise comparisons), followed by abundance (nearly a third) and forest structure (almost a fifth). Forest structure was the most sensitive measure to human disturbance – unsurprising given that anthropogenic land-use changes necessitate (often extensive) alterations to forest structure – followed by abundance and richness. The slightly greater sensitivity of richness compared to abundance is consistent with a past global meta-analysis on tropical forests^49^. This pattern is also consistent with past work showing that abundance of generalist species can increase together with a decline in overall species richness following disturbance in tropical forests^6,37,42,80^. Combined with our result of similar responses to land-use changes across taxa, this suggests that forest specialists, rather than any broad taxonomic group, are particularly susceptible to human disturbance.

In addition to our key *a priori* predictors of effect size variation (disturbance intensity, taxa, and biodiversity metric), we *post hoc* explored whether biodiversity responses to land-use change varied across the region’s four biodiversity hotspots (see Figure 1). We did this by investigating whether model performance was improved by adding ‘biodiversity hotspot’ as an additional predictor (factor) to our top-selected model. This geographic predictor was redundant, with differences in biodiversity responses to land-use change being statistically indistinguishable between hotspots (Δ expected log-pointwise predictive density (see material and methods) = -74.6 ± 23.6, which indicates much greater support for the model without hotspots as a predictor). Although the worst effects of land-use change on biodiversity were found in the Philippines, with a mean reduction of biodiversity in disturbed areas of 38% (CI 7-59%), similar magnitudes of effect were found in Sundaland (35%, CI 27-42%) and Indo-Burma (36%, CI 20-48%). The least severe effects were observed in Wallacea, where the mean reduction was only 5%, but the limited number of highly heterogenous observations from this hotspot meant there was huge amount of uncertainty in our estimate (95% CI: 40% decrease-54% increase).

Biodiversity is a multifaceted concept, comprising a range of components including genetic, taxonomic, phylogenetic, and ecological^81^. A limitation of our study is a reliance on simple measures of biodiversity (i.e., species richness, abundance, and forest structure) that cannot fully capture the complexity of biodiversity and its response to land-use change. For example, a previous global analysis found that, on average, human-affected areas lost narrow-ranged species and gained wide-ranged species^42^, indicating that land-use change might generally have consequences not just for species loss, but also for the homogenisation of biota across space. Other global syntheses have shown that while species richness is often unaffected by land-use change, there can nonetheless be significant effects on species turnover and assemblages^43^ that are variable across ecosystems and environmental gradients^82^. For example, while our results for Southeast Asia show that mature plantations and mature secondary forests recover similar levels of abundance and species richness to undisturbed forests, it is likely that the species compositions of these ecological communities are greatly altered^70,83^. We were unable to investigate these types of biodiversity responses with our dataset, but future field studies that move beyond simple measures such as abundance or species richness and report more detailed metrics (and/or provide their raw data) should allow future syntheses to provide a more detailed picture of the impacts on biodiversity from a range of anthropogenic impacts.

Here we have focused on land-use change as a key driver of biodiversity loss, but its synergies with other global-scale drivers of the biodiversity crisis remains relatively unknown^85^. More recently, there have been greater efforts to investigate the combined impacts of climate- and land-use change on terrestrial biota^41,46,85-88^. These studies generally indicate that the combined of effects of these two drivers will be greater than their additive impact alone, with Southeast Asian biodiversity predicted to be at increased risk from land-cover change as a result of the interaction with climate change, caused for example by drying of forests and increases in wildfires^46,86^. Further extensions of our analyses that include potential synergies of land-use change with other important drivers of tropical biodiversity loss including hunting^89,90^, wildlife trade^11^, and forest fragmentation^91^, will be crucial to fully understanding and potentially mitigating the suite of threats to Southeast Asia’s diverse biota.

## Methods

### Literature search

We searched ISI Web of Science and Scopus on 7 January 2020 for peer-reviewed English studies published between 2010 and 2019, and that contained pairwise comparison of biodiversity values in primary or mature secondary forests and forests subject to land use change and human disturbance. A full list of search terms is provided in Supplementary Information Text S1.

After removing duplicates, we screened the titles and abstracts of 5414 articles in the software Rayyan^92^, removing those that were not broadly relevant to our meta-analysis (i.e., the title and abstract did not contain any information on land-use-change or anthropogenic forest disturbance effects on biodiversity in Southeast Asia). This left 414 papers for which we read the full text and applied the inclusion criteria outlined in the PRISMA (Preferred Reporting Items for Systematic Reviews and Meta-Analyses^93^) diagram (Fig. S1). We included studies that: (1) included some measure of biodiversity across multiple sites and that included areas that could be clearly categorised as ‘control’ sites (primary forest, minimally impacted primary forest or mature secondary forest) and ‘disturbed’ comparison sites; (2) reported means, sample sizes and variance measures for biodiversity in both the control and comparison sites, making them suitable for formal meta-analysis; and (3) were located in the focal study region of Southeast Asia (see Figure 1 for map of area defined as Southeast Asia). After applying these criteria, 173 studies remained. We further refined this set of studies during the data-extraction stage (see next section, *Data extraction and refinement*), resulting in final meta-analysis dataset for modelling comprising 831 effect sizes from 101 papers. The full process of study identification through to inclusion is outlined in the PRISMA diagram (Fig. S1), and full list of studies included in the meta-analysis are given in Supplementary Information Text S2.

### Data extraction and refinement

From each paper, we sought to extract the means, standard deviations and sample sizes of biodiversity measures in control and disturbed comparison sites, to enable the calculation of effect sizes (detailed in next section, *Effect size calculation*). For papers where summary statistics were not reported in the main text, but instead presented in figures, we extracted the data using the R package metaDigitise^94^.

For each comparison, we recorded the country and coordinates of the study. We also recorded the broad taxonomic group studied (plant, invertebrate or vertebrate).

We classified each data point as belonging to one of five categories of biodiversity measures: abundance (for example, number of individuals, density, biomass); richness (for example, observed/estimated/rarefied richness at species/genera/family level); forest structure (for example canopy height/openness, basal area); demographics (for example, mortality rate or size); and community (for example, diversity or abundance of different feeding guilds). However, demographic and community measures were excluded prior to data analysis due to low sample sizes for these categories (n = 7 studies). We further excluded studies when the only reported biodiversity measure was an index (e.g., Shannon’s, Simpson’s or Margalef’s) because these indices are typically secondary (derived) measures of abundance and/or richness^95^ and are often non-linear descriptors of biodiversity change (and so are not directly interpretable in the context of a meta-analysis).

We recorded the disturbance type as it was classified by the study authors, forming eight distinct categories: degraded or young secondary forest, selectively logged forest, clear-cut forest, polyculture plantation, monoculture plantation, urban, restoration area or abandoned agriculture, and forest edge or fragment. We further refined these categories based on the GLOBIO3 framework^38^, splitting disturbance types into high intensity (clear-cut forest, monoculture plantation, urbanised area), low intensity (degraded forest or young, secondary forest, selectively logged forest, polyculture plantation, restored forest) and forest edge or fragment. Prior to analysis, we excluded cases where it was not clear what type of disturbance had occurred in the comparison site. We also excluded comparisons where the disturbed site was a forest edge or fragment, as sample size for this category was too low for meaningful meta-analysis (n = 6 studies).

### Effect-size calculation

For each comparison, we calculated Hedges’ *g*^96^, being the difference between control and disturbed group means, standardised using the pooled standard deviation of the two groups^53^. This was calculated using the mean, standard deviation and sample sizes for the pairwise comparisons, using the following equations implemented through the ‘escalc’ function in the R package metafor^97^:

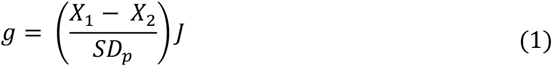

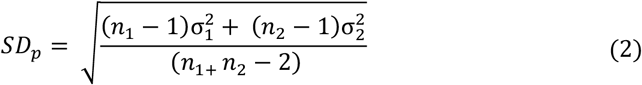

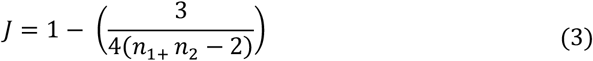

where *X* is the mean, *n* is the sample size, and *σ*^2^ is the variance, for the controls 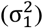 or disturbed 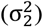 treatment. *J* is a correction factor to remove bias when sample sizes are small^53^. The sampling variance in Hedges’ 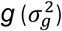 is calculated following^96^ as:

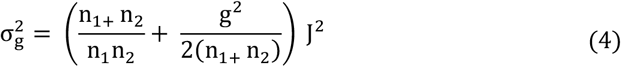

Similar to Cohen’s *d*, Hedges’ *g* expresses the difference in means in terms of standard deviations (making it dimensionless, and therefore allowing comparisons between measurements made in different units), but it is more robust to unequal sampling and small sample sizes^53^. In all cases, we selected an expected direction for the effect size calculation, such that in our analysis, negative effect sizes indicate lower biodiversity in disturbed sites compared to controls. In addition, we calculated log-response ratios^98^ (see Supplementary Information Text S3) from our intercept-only and top-selected model (which contained disturbance type as a predictor), as well as for our analyses of differences between disaggregated land-use-change categories and between biogeographic regions; this allowed us back-transform effect sizes by exponentiating model parameter estimates, thereby expressing changes in biodiversity between control and comparison sites as easy-to-interpret percentage differences.

Most of our 101 papers reported multiple effect sizes, giving us a dataset with 831 total effect sizes. This resulted from studies reporting multiple measures of biodiversity (for example both abundance and richness), multiple comparisons from a single control site to different disturbed sites, and/or multiple measures from the same control-disturbed comparison but across different taxa. We controlled statistically for these non-independent effect sizes in our data analysis (see next section on *Random effects meta-analysis*).

### Random effects meta-analysis and model selection

First, we obtained a weighted mean effect size for the entire dataset, using a Bayesian multi-level meta-analytic model. This mean was obtained by fitting a model with no predictor variables (i.e., an intercept-only model), but included a cluster-level intercept to account explicitly for the non-independence of effect sizes sourced from the same cluster – ‘cluster ID’ is a unique ID given to pairwise comparisons sharing a common control (effectively corresponding to ‘study ID’). This cluster term also partially accounts for spatial correlation of effect sizes that are spatially grouped within each cluster. The intercept-only model is the null submodel of the full location-scale model specified below.

Second, we estimated the amount of heterogeneity present in the dataset. We calculated the statistic *I*^2^ (see Equation 11 for the definition) as an estimate of the proportion of variance in effect size that is due to differences between levels of a random effect (e.g., studies)^99^. *I*^2^ is preferred over other heterogeneity statistics as it is independent of sample size, is easily interpretable and can be partitioned between different random effects^100^. Heterogeneity in ecological and evolutionary datasets is often high, with the mean *I*^2^ from 86 studies in a recent review above 90%^58^.

Third, we fit and compared a set of candidate models to understand which components are most important for predicting heterogenous effects of land-use-change on biodiversity. All the candidate models for our analysis are submodels of the following multilevel Gaussian distribution with modelled location parameter *μ* and modelled scale parameter *σ*:

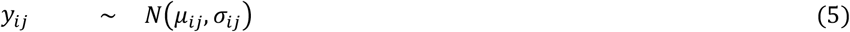

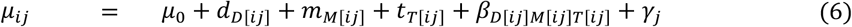

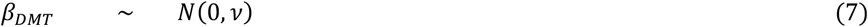

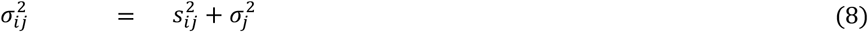

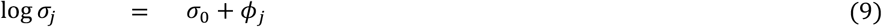

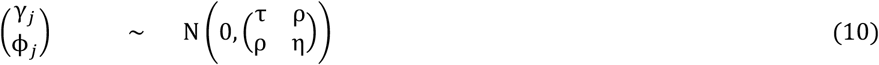

where *y*_*i j*_ is the (standardised) effect size of observation *i* within cluster *j*. For the *μ* −model: *μ*_0_is a global intercept; *d*_*D*_, *m*_*M*_ and *t*_*T*_ are fixed effects for the three (categorical) predictors, disturbance type, biodiversity metric, and taxon, respectively; *β*_*DMT*_ is an interaction-level intercept for all interaction combinations between the levels of three predictors; and *γ*_*i*_ is a cluster-level intercept. The subscripts *D*[*i j*], *M*[*i j*] and *T*[*i j*] indicate the corresponding predictor levels for the datum *y*_*i j*_ and there is no parameter estimate for the first level in each predictor (i.e., a treatment contrast for parameter identifiability). The hyperparameter *ν* provides some regularisation to aid parameter estimation (i.e., model convergence) when interactions *β* are included in the submodel and *τ* is the between-cluster variance of the (cluster-level) intercept estimates. For the *σ*-model: *σ*_0_ is a global intercept, *s*_*i j*_ the study-associated standard errors, and ϕ_*i*_ are cluster level intercepts (*σ*_*j*_ is modelled on the log scale to ensure positivity). The parameter ϕ_*i*_ determines the cluster-level variance and the associated hyperparameter *η* determines the variance of the variance (i.e., the between-cluster variability of the variance estimates). The parameter *ρ* models the covariance of cluster-level intercepts between the location and scale parts of the model. We used weakly informative priors for all parameters (Student-*t* for the intercept, standard deviation, and predictors).

To investigate the role of predictors and their interactions we specified the following set of *µ*-submodels: Null, D, M, T, M+T, D+T, M+D, M+T+D, and Full, where Null includes only global and cluster-level intercepts, the D, M, and T denote the addition of the corresponding fixed effects to the null model (D – disturbance intensity, M – metric, T - taxon), and only the full model includes interactions. To investigate models of differing complexity for the scale parameter, we specified three submodels: *s*_0_(*σ*_*j*_ = 0), *s*_1_(ϕ_*i*_ = 0), and *s*_2_ (the full *σ*-model). Posterior predictive checks and formal model selection showed that the *s*_2_ models performed far better than both the *s*_0_ and *s*_1_ variants (Fig. S2) and so we focus only on results from the *s*_2_ variants in the main text. The full model selection table including the *s*_0_ and *s*_1_ variants is presented in Fig. S3.

To perform model selection, we used 20-fold leave-cluster-out cross validation to estimate the expected log-pointwise predictive density (ELPD) of the marginal model (i.e., predicting to new sites in new clusters)^101^. To select a model for interpretation, we used a modification of the one standard error selection rule^102^ that identifies a parsimonious model by directly accounting for estimation uncertainty in the chosen score^103^. The reported effect and uncertainty estimates were taken from the most parsimonus model selected (the M+D model, Figure 2), yet it is known that post-selection inference on parameter estimates can be biased due to failure to account for model-selection uncertainty^104^. To check the validity of these inferences, we compared the estimates to those of the full model which is known to closely approximate valid post-selection inferences^105^. We found the size of the relevant parameter estimates were virtually identical and led to the same conclusions (e.g., intercept for M+D model = 0.71, 95% CIs 0.50-0.94, intercept for full model = 0.77, 0.48-1.07).

The definition of the *I*^2^ statistic must be slightly adjusted to account for the added complexity of including cluster-level terms in both the location and scale parts of the model^54^. Retaining the usual notion that *I*^2^ is a measure of the proportion of the total variance explained by the between-cluster variance of mean, we take the cluster average of the standard *I*^2^ definition:

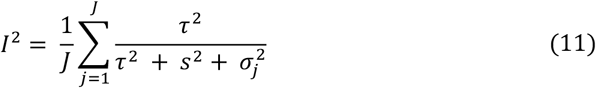

where *j* = 1, …, *J* indexes clusters and 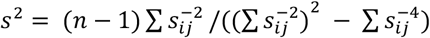 summarises the “typical” within-sample variance^54,106^.

All models were run in the R package brms^107^, each with 4 chains, 4000 sampling iterations and a burn-in of 2000 iterations. We were satisfied with convergence of the marginal Markov chains by a combination of visual inspection of posterior predictive checks, an Rhat statistic under 1.01 and effective sample sizes (bulk and tail) greater than 2000^107^.

### Publication bias

We tested for publication bias (specifically small study bias) statistically via a variant of Egger’s test^108^ that uses the meta-analytic residuals as the response, and precision as the predictor^57^. Publication bias occurs when effect sizes are imbalanced in the literature, typically indicated by an over-representation of large effect sizes with small precision. We found no evidence for bias, with the intercept of Egger’s regression no different from zero (−0.02, -0.4-0.01).

## Acknowledgments

This work was funded by the Australian Research Council grant FL160100101 (awarded to BWB).

## Notes

### Competing Interest Statement

The authors have declared no competing interest.

