## Supplementary Information for "Land use change drives major loss of Southeast Asian biodiversity"

### Supplementary Information for
### Land use change drives major loss of Southeast Asian biodiversity

Thomas Botterill-James[1*], Luke A. Yates[1], Jessie C. Buettel[1,2], Zach Aandahl[1], Barry W. Brook[1,2]

[1]School of Natural Sciences, University of Tasmania; Hobart, 7005, Australia, [2]ARC Centre of Excellence for Australian Biodiversity and Heritage (CABAH)

*Corresponding author: Thomas Botterill-James

**This PDF file includes:**

Supporting text S1 to S3
Figures S1 to S3

**Supplementary Information Text S1: Search terms used to identify relevant papers for meta-analysis (from start 2010-end 2019)**

**Web of Science**

**Advanced search.** TS=(habitat* OR forest* OR land* OR clear-cut* OR disturb* OR plantation* OR fragment*) AND TS=(biodiversity* OR bird* OR mammal* OR amphib* OR reptil* OR vertebrate* OR plants* OR lepidoptera* OR hymenoptera* OR arachnid* OR coleoptera* OR diptera* OR homoptera* OR isoptera* OR invertebrate*) AND TS=(South-east Asia* OR Southeast Asia* OR Brunei* OR Cambodia* OR Indonesia* OR Lao* OR Malaysia* OR Myanmar* OR Burma* OR Philippines* OR Singapore* OR Thailand* OR Vietnam*)

**Scopus**

**Advanced search.** TITLE-ABS-KEY(habitat* OR forest* OR land* OR clear-cut* OR disturb* OR plantation* OR fragment*) AND TITLE-ABS-KEY(biodiversity* OR bird* OR mammal* OR amphib* OR reptil* OR vertebrate* OR plants* OR lepidoptera* OR hymenoptera* OR arachnid* OR coleoptera* OR diptera* OR homoptera* OR isoptera* OR invertebrate*) AND TITLE-ABS-KEY(South-east Asia* OR Southeast Asia* OR Brunei* OR Cambodia* OR Indonesia* OR Lao* OR Malaysia* OR Myanmar* OR Burma* OR Philippines* OR Singapore* OR Thailand* OR Vietnam*) AND PUBYEAR AFT 2009

**Supplementary Information Text S2: Full list of studies included in meta-analysis**


1 C. Agus *et al.,* The effect of tropical peat land-use changes on plant diversity and soil properties. *International Journal of Environmental Science and Technology* **17**, 1703-1712 (2019).

2 J. C. C. Huang, E. L. Rustiati, M. Nusalawo, T. Kingston, Echolocation and roosting ecology determine sensitivity of forest-dependent bats to coffee agriculture. *Biotropica* **51**, 757-768 (2019).

3 E. Yirdaw, A. M. Monge, D. Austin, I. Toure, Recovery of floristic diversity, composition and structure of regrowth forests on fallow lands: implications for conservation and restoration of degraded forest lands in Laos. *New Forests* **50**, doiI; 10.3389/fmicb.2019.00240 1007-1026 (2019).

4 G. Schulz, *et al.,* Changes in trophic groups of protists with conversion of rainforest into rubber and oil palm plantations. *Frontiers in Microbiology* **10**, (2019).

5 R. Koneri, M. J. Nangoy, Butterfly community structure and diversity in Sangihe Islands, North Sulawesi, Indonesia. *Applied Ecology and Environmental Research* **17**, 2501-2517 (2019).

6 J. Jamhuri *et al.,* Selective logging causes the decline of large-sized mammals including those in unlogged patches surrounded by logged and agricultural areas. *Biological Conservation* **227**, 40-47 (2018).

7 T. Bohnert *et al.,* Effects of land-use change on vascular epiphyte diversity in Sumatra (Indonesia). *Biological Conservation* **202**, 20-29 (2016).

8 D. Chellaiah, C. M. Yule, Riparian buffers mitigate impacts of oil palm plantations on aquatic macroinvertebrate community structure in tropical streams of Borneo. *Ecological Indicators* **95**, 53-62 (2018).

9 C. L. Wilkinson, D. C. J. Yeo, H. H. Tan, A. H. Fikri, R. M. Ewers, The availability of freshwater fish resources is maintained across a land-use gradient in Sabah, Borneo. *Aquatic Conservation-Marine and Freshwater Ecosystems* **28**, 1044-1054 (2018).

10 A. Paoletti *et al.,* Amphibian and reptile communities of upland and riparian sites across Indonesian oil palm, rubber and forest. *Global Ecology and Conservation* **16**, e00492 (2018).

11 K. O. Chua *et al.,* Microbial community composition reveals spatial variation and distinctive core microbiome of the weaver ant Oecophylla smaragdina in Malaysia. *Scientific Reports* **8**, 10777 (2018).

12 D. F. R. Cleary Impact of logging on tree, liana and herb assemblages in a Bornean forest. *Journal of Sustainable Forestry* **36**, 806-817 (2017).

13 E. Psomas, S. Holdsworth, P. Eggleton, Ant diversity as a direct and indirect driver of pselaphine rove beetle (Coleoptera: Staphylinidae) functional diversity in tropical rainforests, Sabah, Malaysian Borneo. *Journal of Morphology* **279**, 981-996 (2018).

14 C. L. Wilkinson, D. C. J. Yeo, T. H. Hui, A. H. Fikri, R. M. Ewers, Land-use change is associated with a significant loss of freshwater fish species and functional richness in Sabah, Malaysia. *Biological Conservation* **222**, 164-171 (2018).

15 R. E. J. Gray, R. M. Ewers, M. J. W. Boyle, A. Y. C. Chung, R. J. Gill, Effect of tropical forest disturbance on the competitive interactions within a diverse ant community. *Scientific Reports* **8**, 5131 (2018).

16 T. Changbunjong *et al.,* Species diversity and abundance of Tabanus spp. (Diptera: Tabanidae) in different habitats of Thailand. *Journal of Asia-Pacific Entomology* **21**, 134-139 (2018).

17 A. Kimber, P. Eggleton, Strong but taxon-specific responses of termites and wood-nesting ants to forest regeneration in Borneo. *Biotropica* **50**, 266-273 (2018).

18 M. S. Crane, C. T. Strine, T. K. Knierim, T. Artchawakom, P. Suwanwaree, Herpetofaunal species abundance, richness and diversity in a dry tropical forest and agricultural matrix at the Sakaerat Biosphere reserve, Thailand. *Herpetological Conservation and Biology* **13**, 586-597 (2018).

19 K. Darras *et al.,* Birds of primary and secondary forest and shrub habitats in the peat swamp of Berbak National Park, Sumatra. *F1000Research* **7**, doi:10.12688/f1000research.13996.2. (2018).

20 J. C. Redena-Santos, D. V. Thao, M. Schnittler, N. H. A. Dagamac, The first report of composition and occurrence of myxomycete assemblages in protected and unprotected plantation forests: a comparative study in Thai Nguyen City, Northern Vietnam. *Plant Ecology and Evolution* **151**, 231-240 (2018).

21 A. R. Styring *et al.,* Bird community structure in native forest fragments and Acacia mangium plantations in Borneo. *Wilson Journal of Ornithology* **130**, 112-130 (2018).

22 Z. Rosli, M. Zakaria, M. N. Rajpar, Edge effects on foraging guilds of upperstorey birds in an isolated tropical rainforest of Malaysia. *Journal of Animal and Plant Sciences* **28**, 307-320 (2018).

23 T. H. H. Dao, D. Holscher, Fujian cypress and two other threatened tree species in three conservation zones of a nature reserve in north-western Vietnam. *Forest Ecosystems* **4**, 4-29 (2017).

24 J. A. A. Tangena *et al.,* Diversity of mosquitoes (Diptera: Culicidae) attracted to human subjects in rubber plantations, secondary forests, and villages in Luang Prabang Province, Northern Lao PDR. *Journal of Medical Entomology* **54**, 1589-1604 (2017).

25 T. Bourguignon *et al.,* Ant and termite communities in isolated and continuous forest fragments in Singapore. *Insectes Sociaux* **64**, 505-514 (2017).

26 T. F. Dobert, B. L. Webber, J. B. Sugau, K. J. M. Dickinson, R. K. Didham, Logging increases the functional and phylogenetic dispersion of understorey plant communities in tropical lowland rain forest. *Journal of Ecology* **105**, 1235-1245 (2017).

27 S. N. Shuhada, S. Salim, F. Nobilly, A. Zubaid, B. Azhar, Logged peat swamp forest supports greater macrofungal biodiversity than large-scale oil palm plantations and smallholdings. *Ecology and Evolution* **7**, 7187-7200 (2017).

28 K. Rembold, H. Mangopo, S. Tjitrosoedirdjo, H. Kreft, Plant diversity, forest dependency, and alien plant invasions in tropical agricultural landscapes. *Biological Conservation* **213**, 234-242 (2017).

29 C. C. P. Cosset, D. P. Edwards, The effects of restoring logged tropical forests on avian phylogenetic and functional diversity. *Ecological Applications* **27**, 1932-1945 (2017).

30 J. Wills, J. Herbohn, M. O. M. Moreno, M. S. Avela, J. Firn, Next-generation tropical forests: reforestation type affects recruitment of species and functional diversity in a human-dominated landscape. *Journal of Applied Ecology* **54**, 772-783 (2017).

31 O. R. Wearn *et al.,* Mammalian species abundance across a gradient of tropical land-use intensity: A hierarchical multi-species modelling approach. *Biological Conservation* **212**, 162-171 (2017).

32 A. M. Tulod, J. V. Casas, R. A. Marin, J. A. B. Ejoc, Diversity of native woody regeneration in exotic tree plantations and natural forest in Southern Philippines. *Forest Science and Technology* **13**, 31-40 (2017).

33 K. S. Masum, A. Mansor, S. A. M. Sah, H. S. Lim, M. K. Hossain, Effect of differential forest management on biodiversity in a tropical hill forest of Malaysia and implications for conservation. *Biodiversity and Conservation* **26**, 1569-1586 (2017).

34 B. Klarner *et al.,* Trophic niches, diversity and community composition of invertebrate top predators (Chilopoda) as affected by conversion of tropical lowland rainforest in Sumatra (Indonesia). *Plos One* **12**, e0180915 (2017).

35 S. Hosoishi, Y. Hashimoto, S. H. Park, S. Yamane, K. Ogata, A comparison of ground-dwelling and arboreal ant assemblages (Hymenoptera: Formicidae) in lowland forests of Cambodia. *Raffles Bulletin of Zoology* **65**, 416-425 (2017).

36 R. Koneri, M. J. Nangoy, M. J. The distribution and diversity of spiders (Arachnida: Arane), Sahendaruman Mountain, Sangihe Islands, North Sulawesi, Indonesia. *Applied Ecology and Environmental Research* **15**, 797-808 (2017).

37 W. Marthy, Y. Clough, T. Tscharntke, Assessing the potential for avifauna recovery in degraded forests in Indonesia. *Raffles Bulletin of Zoology* **65**, 35-48 (2017).

38 F. A. Edwards *et al.,* The impact of logging roads on dung beetle assemblages in a tropical rainforest reserve. *Biological Conservation* **205**, 85-92 (2017).

39 S. A. Mukul, J. Herbohn, J. Firn, Co-benefits of biodiversity and carbon sequestration from regenerating secondary forests in the Philippine uplands: implications for forest landscape restoration. *Biotropica* **48**, 882-889 (2017).

40 H. Bernard *et al.,* Species richness and distribution of primates in disturbed and converted forest landscapes in Northern Borneo. *Tropical Conservation Science* **9**, doi: 10.1177/1940082916680104 (2016).

41 D. Kerfahi, B. M. Tripathi, K. Dong, R. Go, J. M. Adams, Rainforest conversion to rubber plantation may not result in lower soil diversity of bacteria, fungi, and nematodes. *Microbial Ecology* **72**, 359-371 (2016).

42 O. R Wearn, C. Carbone, J. M. Rowcliffe, H. Bernard, R. M. Ewers, Grain-dependent responses of mammalian diversity to land use and the implications for conservation set-aside. *Ecological Applications* **26**, 1409-1420 (2016).

43 W. E. Prabowo *et al.,* Bird responses to lowland rainforest conversion in Sumatran smallholder landscapes, Indonesia. *Plos One* **11**, e0154876 (2016).

44 B. M. Tripathi *et al.,* Distinctive tropical forest variants have unique soil microbial communities, but not always low microbial diversity. *Frontiers in Microbiology* **7**, doi: 10.3389/fmicb.2016.00376 (2016).

45 A. Magrach *et al.,* Selective logging in tropical forests decreases the robustness of liana - tree interaction networks to the loss of host tree species. *Proceedings of the Royal Society B-Biological Sciences* **283**, doi:10.1098/rspb.2015.3008 (2016).

46 T. Yamada, S. Yoshida, T. Hosaka, T. Okuda, Logging residues conserve small mammalian diversity in a Malaysian production forest. *Biological Conservation* **194**, 100-104 (2016).

47 J. H. Moore, S. Sittimongkol, A. Campos-Arceiz, T. Sumpah, M. P. Eichhorn, Fruit gardens enhance mammal diversity and biomass in a Southeast Asian rainforest. *Biological Conservation* **194**, 132-138 (2016).

48 Y. Nurulita, E. M. Adetutu, H. Gunawan, D. Zul, A. S. Ball, Restoration of tropical peat soils: The application of soil microbiology for monitoring the success of the restoration process. *Agriculture Ecosystems & Environment* **216**, 293-303 (2016).

49 C. O. Delang, X. Weiyi, B. Brooke, K. P. Chun, The effect of fallow period length on the abundance and diversity of usable plant assemblages in shifting cultivation system (swidden agriculture) in northern Laos. *Polish Journal of Ecology* **64**, 350-356 (2016).

50 S. Mumme, M. Jochum, U. Brose, N. F. Haneda, A. D. Barnes, Functional diversity and stability of litter-invertebrate communities following land-use change in Sumatra, Indonesia. *Biological Conservation* **191**, 750-758 (2015).

51 X. Giam *et al.,* Mitigating the impact of oil-palm monoculture on freshwater fishes in Southeast Asia. *Conservation Biology* **29**, 1357-1367 (2015).

52 O. Konopik, I. Steffan-Dewenter, T. U. Grafe, Effects of logging and oil palm expansion on stream frog communities on Borneo, Southeast Asia. *Biotropica* **47**, 636-643 (2015).

53 W. Y. Wang, W. A. Foster, The effects of forest conversion to oil palm on ground-foraging ant communities depend on beta diversity and sampling grain. *Ecology and Evolution* **5**, 3159-3170 (2015).

54 A. Ueda *et al.,* Effect of habitat transformation from grassland to Acacia mangium plantation on dung beetle assemblage in East Kalimantan, Indonesia. *Journal of Insect Conservation* **19**, 765-780 (2015).

55 V. Krashevska, B. Klarner, R. Widyastuti, M. Maraun, S. Scheu, Impact of tropical lowland rainforest conversion into rubber and oil palm plantations on soil microbial communities. *Biology and Fertility of Soils* **51**, 697-705 (2015).

56 R. M. Ewers *et al.,* Logging cuts the functional importance of invertebrates in tropical rainforest. *Nature Communications* **6**, 6836, (2015).

57 Y. Nurulita *et al.,* The assessment of the impact of oil palm and rubber plantations on the biotic and abiotic properties of tropical peat swamp soil in Indonesia. *International Journal of Agricultural Sustainability* **13**, 150-166 (2015).

58 W. Asfiya, L. Lach, J. D. Majer, B. Heterick, R. K. Didham, Intensive agroforestry practices negatively affect ant (Hymenoptera: Formicidae) diversity and composition in southeast Sulawesi, Indonesia. *Asian Myrmecology* **7**, 87-105 (2015).

59 M. Singh, Y. Malhi, S. A. Bhagwat, Aboveground biomass and tree diversity of riparian zones in an oil-palm dominated mixed landscape in Borneo. *Journal of Tropical Forest Science* **27**, 227-239 (2015).

60 O. Konopik, C. L. Gray, T. U. Grafe, I. Steffan-Dewenter, T. M. Fayle, From rainforest to oil palm plantations: Shifts in predator population and prey communities, but resistant interactions. *Global Ecology and Conservation* **2**, 385-394 (2014).

61 D. P. Edwards *et al.,* Selective-logging and oil palm: multitaxon impacts, biodiversity indicators, and trade-offs for conservation planning. *Ecological Applications* **24**, 2029-2049 (2014).

62 D. Kerfahi, B. M. Tripathi, J. Lee, D. P. Edwards, J. M. Adams, The impact of selective-logging and forest clearance for oil palm on fungal communities in Borneo. *Plos One* **9**, e111525 (2014).

63 M. Hasegawa *et al.,* The effects of reduced-impact logging practices on soil animal communities in the Deramakot Forest Reserve in Borneo. *Applied Soil Ecology* **83**, 13-21 (2014).

64 S. H. Luke, T. M. Fayle, P. Eggleton, E. C. Turner, R. G. Davies, Functional structure of ant and termite assemblages in old growth forest, logged forest and oil palm plantation in Malaysian Borneo. *Biodiversity and Conservation* **23**, 2817-2832 (2014).

65 Arbainsyah, H. H. Iongh, W. Kustiawan, G. R. de Snoo, Structure, composition and diversity of plant communities in FSC-certified, selectively logged forests of different ages compared to primary rain forest. *Biodiversity and Conservation* **23**, 2445-2472 (2014).

66 C. L. Gray, E. M. Slade, D. J., Mann, O. T. Lewis, Do riparian reserves support dung beetle biodiversity and ecosystem services in oil palm- dominated tropical landscapes? *Ecology and Evolution* **4**, 1049-1060 (2014).

67 F. A. Edwards *et al.,* Does logging and forest conversion to oil palm agriculture alter functional diversity in a biodiversity hotspot? *Animal Conservation* **17**, 163-173 (2014).

68 J. A Hanis, A. Abu Hassan, A. T. Nurita, M. R. C. Salmah, Community structure of termites in a hill dipterocarp forest of Belum-Temengor Forest Complex, Malaysia: emergence of pest species. *Raffles Bulletin of Zoology* **62**, 3-11 (2014).

69 A. Junsongduang, H. Balslev, A. Jampeetong, A. Inta, P. Wangpakapattanawong, Woody Plant Diversity in Sacred Forests and Fallows in Chiang Mai, Thailand. *Chiang Mai Journal of Science* **41**, 1132-1149 (2014).

70 J. M. Lucey *et al.,* Tropical forest fragments contribute to species richness in adjacent oil palm plantations. *Biological Conservation* **169**, 268-276 (2014).

71 D. Muhamad, S. Okubo, T. Miyashita, Parikesit, K. Takeuchi, Effects of habitat type, vegetation structure, and proximity to forests on bird species richness in a forest-agricultural landscape of West Java, Indonesia. *Agroforestry Systems* **87**, 1247-1260 (2013).

72 P. Thongsripong *et al.,* Mosquito vector diversity across habitats in central Thailand endemic for dengue and other arthropod-borne diseases. *Plos Neglected Tropical Diseases* **7**, e2507 (2013).

73 Y.. Tokumoto, T. Itioka, T. Ohkubo, O. Tadauchi, M. Nakagawa, Assemblage of flower visitors to Dillenia suffruticosa and possible negative effects of disturbances in Sarawak, Malaysia. *Entomological Science* **16**, 341-351 (2013).

74 R. L. Kitching, L. A. Ashton, A. Nakamura, T. Whitaker, C. V. Khen, Distance-driven species turnover in Bornean rainforests: homogeneity and heterogeneity in primary and post-logging forests. *Ecography* **36**, 675-682 (2013).

75 A. Faruk, D. Belabut, N. Ahmad, R. J. Knell, T. W. J. Garner, Effects of oil-palm plantations on diversity of tropical anurans. *Conservation Biology* **27**, 615-624 (2012).

76 F. A. Edwards, D. P. Edwards, K. C. Hamer, R. G. Davies, Impacts of logging and conversion of rainforest to oil palm on the functional diversity of birds in Sundaland. *Ibis* **155**, 313-326 (2013).

77 C. C. Van, L. David, H. Marc, Simple plantations have the potential to enhance biodiversity in degraded areas of Tam Dao National Park, Vietnam. *Natural Areas Journal* **33**, 139-147 (2013).

78 M. Yoshima, Y. Takematsu, A. Yoneyama, M. Nakagawa, Recovery of litter and soil invertebrate communities following swidden cultivation in Sarawak, Malaysia. *Raffles Bulletin of Zoology* **61**, 767-777 (2013).

79 M. Nakagawa *et al.,* Tree community structure, dynamics, and diversity partitioning in a Bornean tropical forested landscape. *Biodiversity and Conservation* **22**, 127-140 (2013).

80 N. Imai, *et al.,* Effects of selective logging on tree species diversity and composition of Bornean tropical rain forests at different spatial scales. *Plant Ecology* **213**, 1413-1424 (2012).

81 V. T. Thinh, P. F. Doherty, K. P. Huyvaert, Avian conservation value of pine plantation forests in northern Vietnam. *Bird Conservation International* **22**, 193-204 (2012).

82 J. M. Lucey, J. K. Hill, Spillover of insects from rain forest into adjacent oil palm plantations. *Biotropica* **44**, 368-377 (2012).

83 D. P. Edwards *et al.,* Reduced-impact logging and biodiversity conservation: a case study from Borneo. *Ecological Applications* **22**, 561-571 (2012).

84 P. Addo-Fordjour, Z. B. Rahmad, A. M. S. Shahrul, A. M. S. Effects of human disturbance on liana community diversity and structure in a tropical rainforest, Malaysia: implication for conservation. *Journal of Plant Ecology* **5**, 391-399 (2012).

85 B. Fisher *et al.,* Cost-effective conservation: calculating biodiversity and logging trade-offs in Southeast Asia. *Conservation Letters* **4**, 443-450 (2011).

86 P. Woodcock *et al.,* The conservation value of South East Asia's highly degraded forests: evidence from leaf-litter ants. *Philosophical Transactions of the Royal Society B-Biological Sciences* **366**, 3256-3264 (2011).

87 M. R. C. Posa, Peat swamp forest avifauna of Central Kalimantan, Indonesia: Effects of habitat loss and degradation. *Biological Conservation* **144**, 2548-2556 (2011).

88 N. Z. Htun, N. Mizoue, S. Yoshida, Tree species composition and diversity at different levels of disturbance in Popa Mountain Park, Myanmar. *Biotropica* **43**, 597-603 (2011).

89 P. Phommexay, C. Satasook, P. Bates, M. Pearch, S. Bumrungsri, The impact of rubber plantations on the diversity and activity of understorey insectivorous bats in southern Thailand. *Biodiversity and Conservation* **20**, 1441-1456 (2011).

90 A. R. Styring *et al.,* Bird community assembly in Bornean industrial tree plantations: Effects of forest age and structure. *Forest Ecology and Management* **261**, 531-544 (2011).

91 K. Kishimoto-Yamada, T. Itioka, M. Nakagawa, K. Momose, T. Nakashizuka, Phytophagous Scarabaeid diversity in swidden cultivation landscapes in Sarawak, Malaysia. *Raffles Bulletin of Zoology* **59**, 285-293 (2011).

92 F. H. Sheldon, A. R. Styring, Bird diversity differs between industrial tree plantations on Borneo: Implications ofr conservation planning. *Raffles Bulletin of Zoology* **59**, 295-309 (2011).

93 P. Hoehn, I. Steffan-Dewenter, T. Tscharntke, Relative contribution of agroforestry, rainforest and openland to local and regional bee diversity. *Biodiversity and Conservation* **19**, 2189-2200 (2010).

94 T. M. Fayle, *et al.,* Oil palm expansion into rain forest greatly reduces ant biodiversity in canopy, epiphytes and leaf-litter. *Basic and Applied Ecology* **11**, 337-345 (2010).

95 S. Kitamura, S. Thong-Aree, S. Madsri, P. Poonswad, Mammal diversity and conservation in a small isolated forest of Southern Thailand. *Raffles Bulletin of Zoology* **58**, 145-156 (2010).

96 T. Okuda *et al.,* Canopy height recovery after selective logging in a lowland tropical rain forest. *Forest Ecology and Management* **442**, 117-123 (2010).

97 K. C. Tanalgo, M. Achondo, A. C. Hughes, Small things matter: the value of rapid biodiversity surveys to understanding local bird diversity patterns in Southcentral Mindanao, Philippines. *Tropical Conservation Science* **12**, doi:10.1177/1940082919869482 (2010).

98 A. Jain, F. K. S. Lim, E. L. Webb, Species-habitat relationships and ecological correlates of butterfly abundance in a transformed tropical landscape. *Biotropica* **49**, 355-364 (2010).

99 A. S. Nurinsiyah, H. Fauzia, C. Hennig, B. Hausdorf, Native and introduced land snail species as ecological indicators in different land use types in Java. *Ecological Indicators* **70**, 557-565 (2010).

100 G. Stride *et al.,* Contrasting patterns of local richness of seedlings, saplings, and trees may have implications for regeneration in rainforest remnants. *Biotropica* **50**, 889-897 (2018).

101 D. F. R. Cleary, K. A. O. Eichhorn, Variation in the composition and diversity of ground-layer herbs and shrubs in unburnt and burnt landscapes. *Journal of Tropical Ecology* **34**, 243-256 (2018).

**Supplementary Information Text S3: Log response ratio calculation**

We calculated log response ratios from our intercept-only and top-selected model, as well as for our analyses of differences between disaggregated land use change categories and between biogeographic regions, so we could exponentiate model parameters to express biodiversity change between control and comparison sites as easy to interpret percentage differences. For each comparison, we calculated the log response ratio, being the ratio of means between control and disturbed group means. This was calculated using the mean, standard deviation and sample sizes for the pairwise comparisons, using the following equations implemented through the escalc function in the R package *metafor*:

$$lnR = \ln(R) = \ln\left(\frac{\bar{X}_1}{\bar{X}_2}\right) = \ln(\bar{X}_1) - \ln(\bar{X}_2) \tag{1}$$

Where $lnR$ is the log response ratio, $\bar{X}_1$ is the mean of the control group and $\bar{X}_2$ is the mean of the disturbed group. The variance of the response ratio is approximately:

$$V_{lnR} = \left(\frac{1}{n_1(\bar{X}_1)^2} + \frac{1}{n_2(\bar{X}_2)^2}\right) \tag{2}$$

Where $S^2{}_{pooled}$ is the pooled standard deviation:

$$SD_p = \sqrt{\frac{(n_1 - 1)\sigma_1^2 + (n_2 - 1)\sigma_2^2}{(n_{1+} n_2 - 2)}} \tag{3}$$

The approximate standard error – used to weight each effect size in the analysis – is:

$$SE_{lnR=\sqrt{V_{lnR}}} \tag{4}$$

In all cases, we selected an expected direction for the effect size calculation, such that in our analysis, negative effect sizes indicate lower biodiversity in disturbed sites compared to controls.

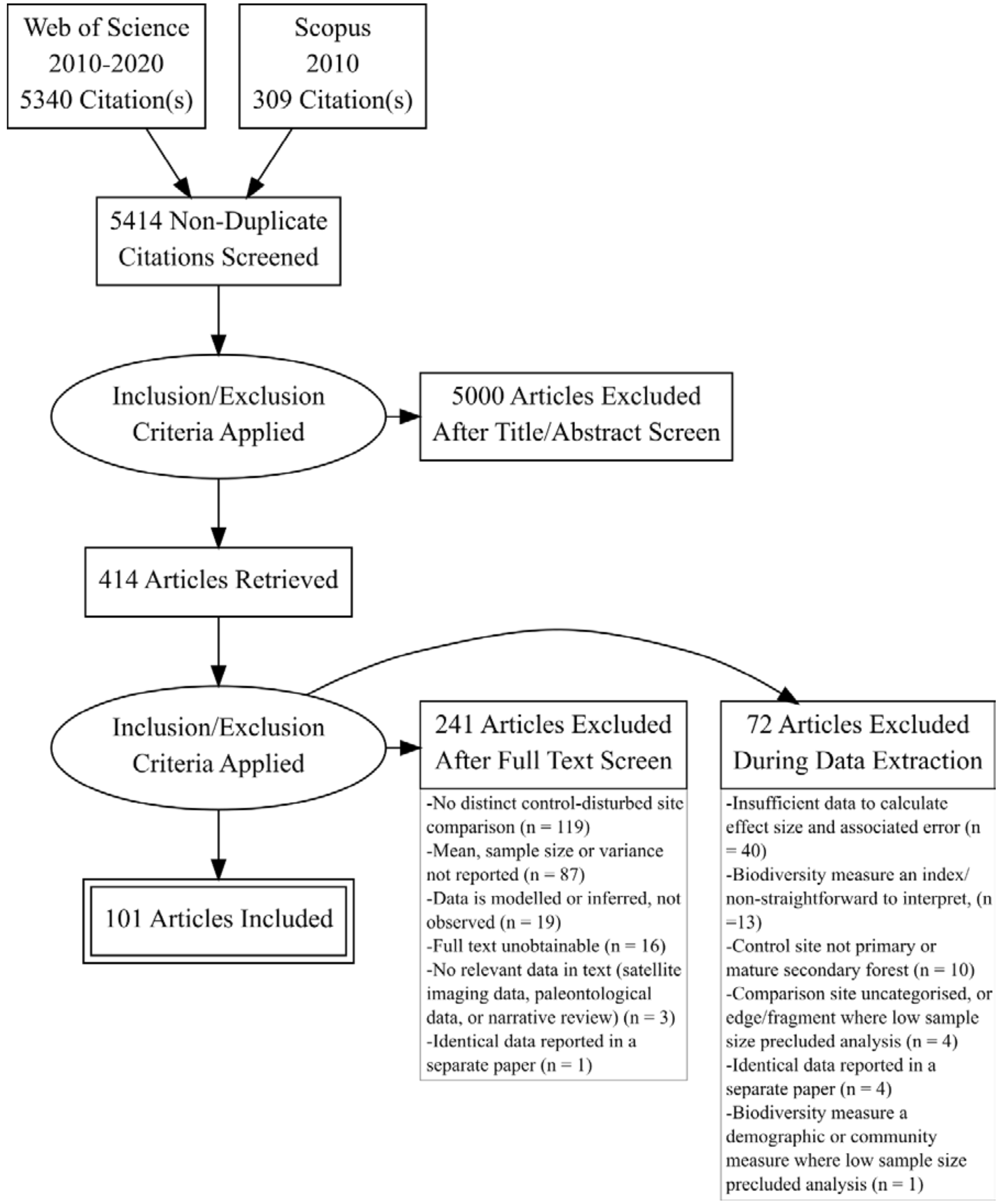


**Fig. S1.** PRISMA diagram. Flow of inclusion and exclusion of studies identified during the literature search and data extraction process, presented as an adapted PRISMA diagram with number of published papers in brackets.

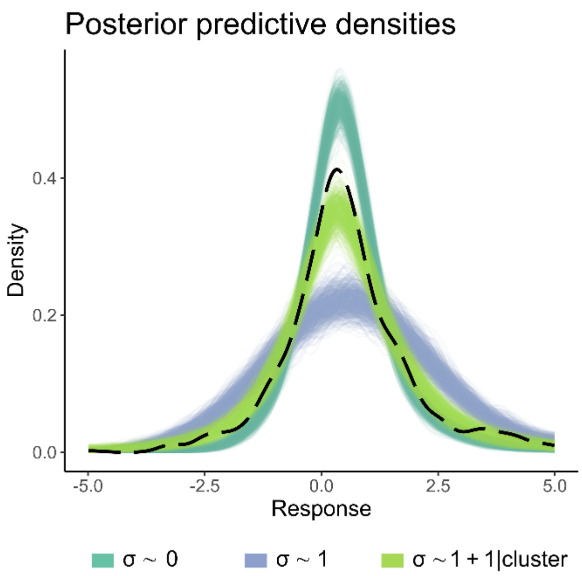


**Fig. S2.** Predictive checks showing simulated data generated using parameters drawn from the posterior distribution of the best model, which contained disturbance intensity and metric type as predictors. These checks illustrate the differing ability of three variants of the selected disturbance-intensity + metric model to produce data comparable to the observed data (dashed line). The variants are characterised by increasing complexity in the modelled structure of the scale parameter: $\sigma \sim 0$ (study associated standard errors only), $\sigma \sim 1$ (inclusion of a global residual error term), and $\sigma \sim 1 + 1|cluster$ (inclusion of a global term plus study-level terms). The latter generates the most plausible data, which together with formal model comparison (see **Fig. S3**), validates its selection as the preferred model for inference in this study. Results were qualitatively similar when varying the scale parameter for the null and full models.

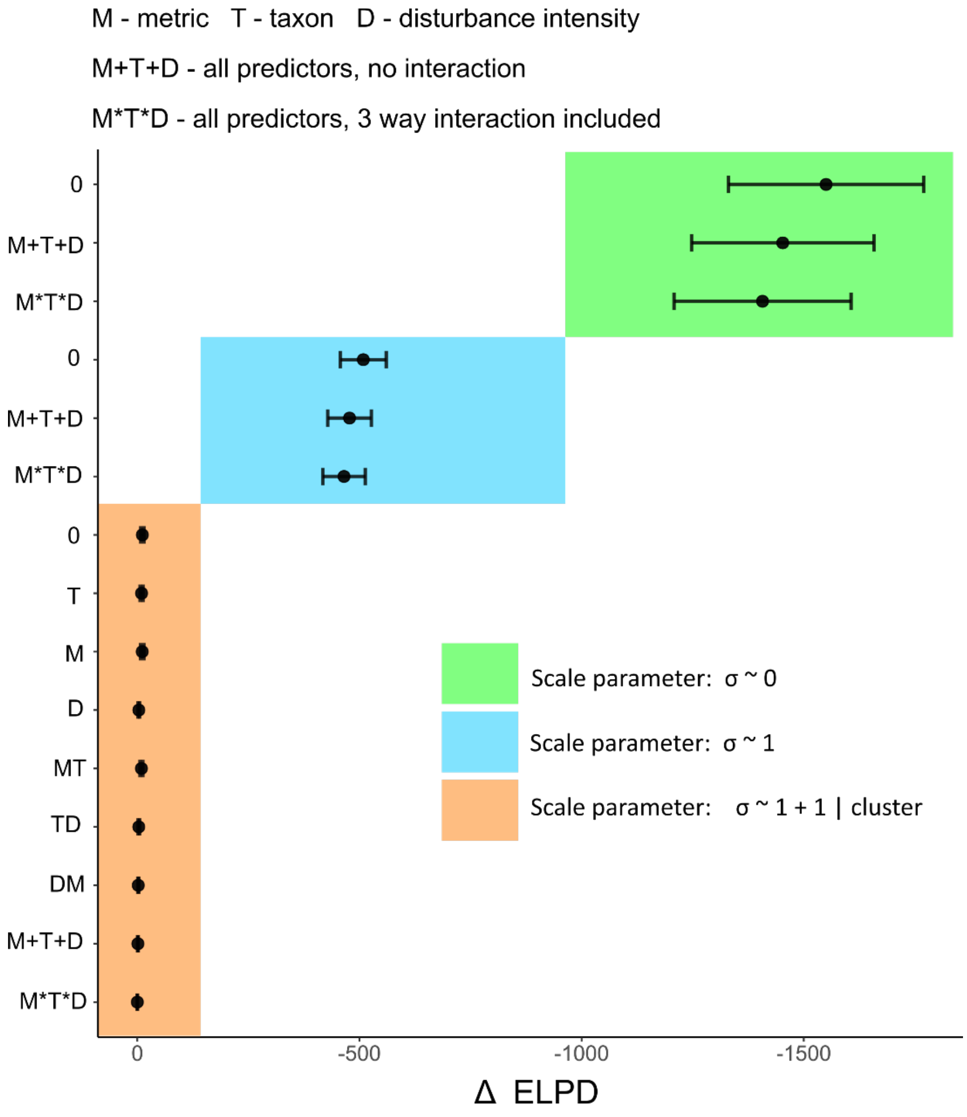


**Fig. S3.** Relative performance of models of biodiveristy metric, taxon and disturbance intensity effects, and their interactions, on biodiversity loss in Southeast Asia. Out-of-sample model performance is estimated here using using leave-one-group(cluster)-out 20-fold cross validation. Models are grouped on the y-axis by increasing complexity in the modelled structure of the scale parameter: σ ~ 0 (study associated standard errors only), σ ~ 1 (inclusion of a global residual error term), and σ ~ 1 + 1|study ID (inclusion of a global term plus study-level terms). Model performance is quantified using expected log-pointwise predictive density (ELPD), expressed here as the relative estimates (Δ ELPD) with respect to the best-performing model. The error bars depict one standard error of the estimated differences. The group of models with the most complex scale parameter perform far better than the other model variants, validating their selection as focus for the manuscript.
